# Sequencing, Chromosome-scale Assembly, and Annotation of the Genome of the Halophilic Nanoflagellate *Halocafeteria seosinensis*

**DOI:** 10.64898/2026.06.25.734631

**Authors:** Lucie Gallot-Lavallée, Ronie Haro, Jon Jerlström-Hultqvist, Dmytro Tymoshenko, Andrew J. Roger, John M. Archibald

**Affiliations:** Department of Biochemistry & Molecular Biology, Dalhousie University, Halifax, Nova Scotia, Canada; Institute for Comparative Genomics, Dalhousie University, Halifax, Nova Scotia, Canada; Department of Cell and Molecular Biology, Uppsala Universitet, Uppsala, Sweden; Department of Biology, Dalhousie University, Halifax, Nova Scotia, Canada

**Author notes:** Corresponding author: J.M. Archibald.

**Keywords:** Extremophile, Halophilic, Hypersaline, Long-read sequencing, Oxford Nanopore, Genome assembly, *Halocafeteria*

## Abstract

Compared with bacterial and archaeal extremophiles, single-celled eukaryotes living in extreme habitats are understudied and underrepresented in genomic databases. An exception is the obligately halophilic stramenopile *Halocafeteria seosinensis* strain EHF34. A transcriptome-focused analysis of this extremophilic protist*s* revealed the importance of organic osmolyte regulation and transport in its adaptation to hypersaline environments. However, genomic resources for *H. seosinensis* are currently limited to a highly fragmented assembly generated by short-read sequencing, which has hindered further investigation of the genome biology and evolution of this fascinating organism. Here, we used long-read Oxford Nanopore sequencing to generate a highly contiguous, chromosome-scale genome assembly for *H. seosinensis*. The assembly is 38.8 megabase pairs (Mbp) in size and contains 60 nuclear contigs, making it the most contiguous genome for a member of the order Bicosoecida. Approximately 19% of the genome is comprised of transposable elements. Of the 11,684 predicted protein-coding genes, many appear to be associated with DNA mobility-related functions, and several may be linked to adaptation to a hypersaline environment. Analysis of the *H. seosinensis* long-read genome assembly presented herein will facilitate our understanding of the ways in which protists have adapted to extreme environments.

**Significance:** *Halocafeteria seosinensis* is an extremophilic protist adapted to hypersaline environments. Previous analyses of a transcriptome and short-read draft genome assembly for this organism provided insights into the molecular mechanisms underlying osmotic regulation, which facilitate its adaptation to high-salt conditions. However, the lack of contiguity and quality of the draft assembly prevented the characterization of complex genomic regions, including transposable elements and viral insertions, as well as genomic comparisons with related species. Here we present a highly contiguous, chromosome-scale genome assembly for *H. seosinensis* that enables accurate gene prediction, detailed analysis of repeat content, and comparative genomic analysis. This long-read genome assembly will serve as a valuable resource for studying one of the few tractable halophilic protists sequenced to date.

## Introduction

The study of eukaryotic genomes has historically focused on model plant and animal species, fungi, and protist pathogens of biomedical and agricultural significance. The diversity of unicellular eukaryotes has remained largely unexplored (Schoenle et al., 2025; Sibbald and Archibald, 2017). This includes extremophilic protists that are far more prevalent in nature than previously believed (Rappaport and Oliverio, 2023) and remain understudied relative to prokaryotic extremophiles (Rappaport and Oliverio, 2024). This knowledge gap is primarily due to the challenges of culturing protists with unknown nutrient requirements and dependency on cobionts (Rappaport and Oliverio, 2023), which impedes the generation of sufficient biomass for high-throughput sequencing. Despite these obstacles, notable progress has been made in sequencing the genomes of select extremophilic protists, such as the thermoacidophilic red alga *Galdieria sulphuraria* (Schönknecht et al., 2013) and the halophilic stramenopile *Halocafeteria seosinensis* (Harding et al., 2016). These advancements have provided valuable insights into the biochemical and cellular adaptations of these organisms to extreme environments.

*H. seosinensis* is a tiny (∼4 μm long) halophilic nanoflagellate originally isolated from solar salterns with salinity levels around 300‰ (Park et al., 2006). The organism naturally grazes on prokaryotes and, although details of its reproduction and life cycle remain scarce, it thrives in high-salinity environments by accumulating organic osmolytes, such as myoinositol, to regulate osmotic balance (Harding et al., 2016). The organism’s highly hydrophilic cytoplasmic proteome is predicted to ensure protein solubility and functionality within its saline intracellular environment (Harding et al., 2016). Within the stramenopiles, the genomic landscape of bicosoecids to which *H. seosinensis* belongs remains poorly characterized, with *Cafeteria burkhardae* (formerly *roenbergensis*) being the only representative with a high-quality, well-described genome (Hackl et al., 2020). Remarkably, analysis of the *C. burkhardae* genome revealed the presence of a thriving community of endogenized virophages (i.e., Mavirus or endogenous mavirus-like elements (EMALEs)) (Hackl et al., 2021). *C*. *burkhardae* is also naturally infected by the giant virus CroV (family *Mimiviridae*) (Fischer et al., 2010). These findings suggest that viruses may significantly shape the genomes of bicosoecids, making this lineage an interesting one in which to study protist-virus interactions.

Although the initial draft genome assembly of *H. seosinensis* offered valuable insights into the molecular mechanisms underlying the organism’s adaptation to hypersaline environments (Harding et al., 2016), it was generated using short-read sequencing technology. The lack of contiguity has restricted further exploration of key genomic features, such as ploidy, repeat content, gene duplications, and the presence and diversity of endogenous viral elements. Here we present a chromosomal-scale genome assembly of *H. seosinensis* strain EHF34, generated using Oxford Nanopore Technologies long-read sequencing. This assembly represents the most complete and contiguous genome assembly presently available for bicosoecids, opening new avenues for comparative genomics and studying the biology of extremophilic protists and the viruses that infect them.

## Results and Discussion

### Genome assembly and repeat content

We generated approximately 2.7 million nanopore reads (∼20.8 gigabase pairs in total), with a read N50 of 13.4 kilobase pairs (Kbp). Approximately 32% of the reads were inferred to be contaminants (mostly prokaryotic), leaving ∼1.8 million reads for reassembly. These reads were assembled into 60 nuclear contigs with ∼235× coverage (Fig. 1a, Supplementary Fig. S1), plus a 44 Kbp contig corresponding to the circular-mapping mitochondrial genome (Supplementary Fig. S2b). 48 of the 60 nuclear contigs were found to have telomeric repeats (TAACCCT^n^) on one end (Fig. 1a), while four contigs (contig_233, 34, 153, and 4) contained obvious telomeres at both ends, consistent with them representing complete or near-complete chromosomes (Fig. 1a). Approximately 92% of the assembly was contained within 50 large contigs totalling 38.6 Mbp (Supplementary Fig. S1a), indicating that the assembly size is likely close to the actual genome size.

**Fig. 1.**
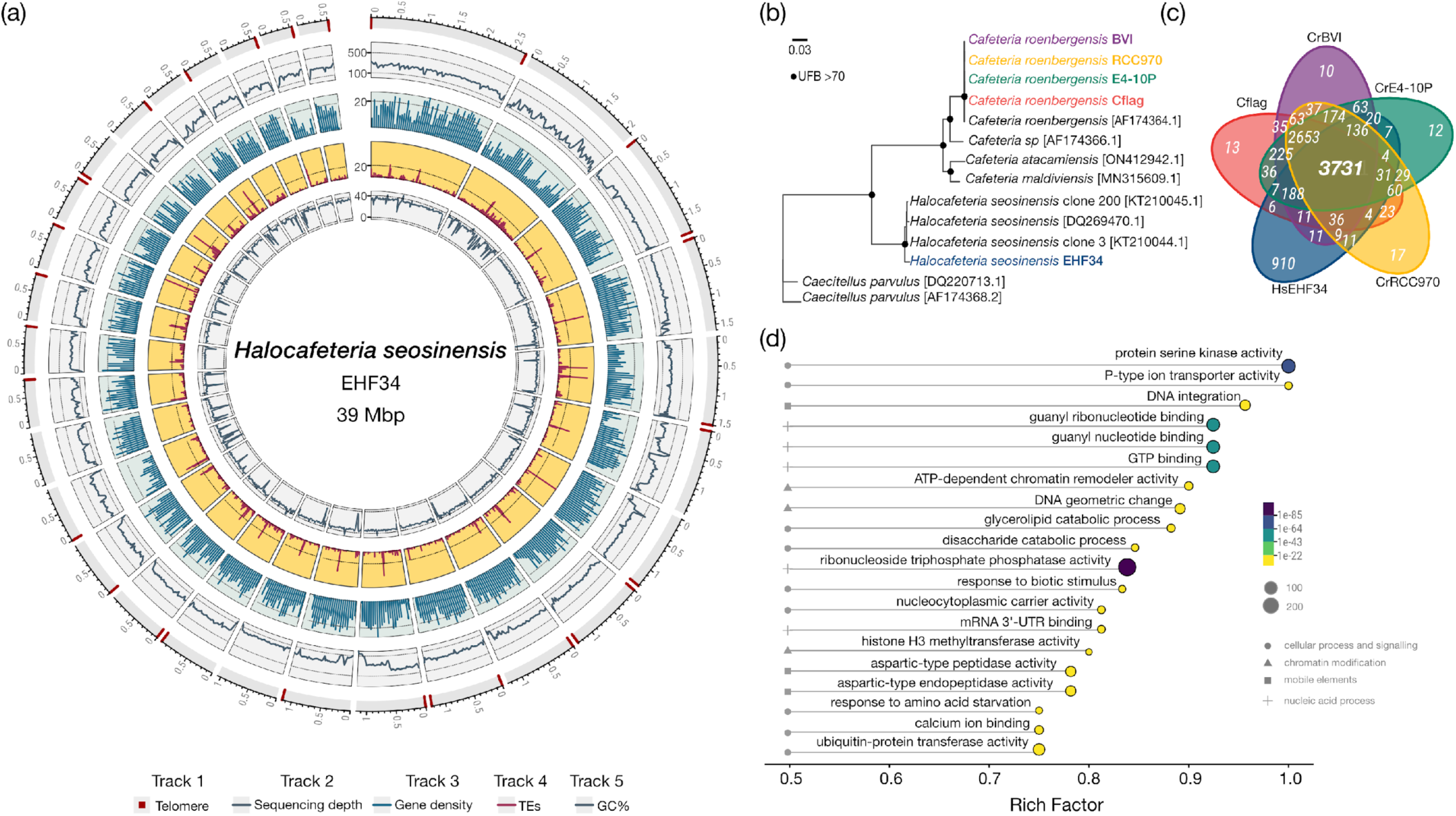
Key features of the *Halocafeteria seosinensis* nuclear genome assembly and comparative genomic analysis with *Cafeteria burkhardae* strains. a) Circos plot of the *H. seosinensis* EHF34 assembly. The outermost track (Track 1) corresponds to contigs > 500 Kbp in size; telomeric repeats on one or both ends are shown in red. Tracks 2 to 5 are described in the legend. b) Maximum likelihood phylogenetic tree showing relationships between select *Cafeteria* spp. strains, *Halocafeteria*, and *Caecitellus* species reconstructed using 18S rRNA gene sequences (∼1.6 Kbp). NCBI access IDs are shown in brackets. The scale bar shows the inferred number of nucleotide substitutions per site. UFB: ultrafast bootstrap. c) Number of orthogroups shared by the four strains of *C. burkhardae* and *H. seosinensis* (HsEHF34). d) Enrichment analysis of the unique gene fraction of *H. seosinensis* (910 orthogroups) showing the 20 most enriched GO categories. Categories are sorted according to Rich Factor.

The completeness and contiguity of the *H. seosinensis* genome assembly presented here represents a substantial improvement over its short-read-based predecessor (Harding et al., 2016), and the assemblies of four *C. burkhardae* strains (Hackl et al., 2020) (Table 1). The relative completeness of these assemblies was assessed using BUSCO odb12 (Tegenfeldt et al., 2025). The *H. seosinensis* long-read assembly was found to contain 82% of the stramenopile BUSCO orthologs and 78% of the eukaryotic orthologs. Meanwhile, the *C. burkhardae* assemblies recovered 87% and 65% BUSCO completeness, respectively. This discrepancy may be due to the absence of *Halocafeteria*-specific orthologs in the Odb12 stramenopile dataset. Completeness of the short-read and long-read-based assemblies of *H. seosinensis* is similar, despite the greater contiguity of the latter (Table 1). The *H. seosinensis* assembly presented herein has a total length of 38.89 Mbp with an N50 of 1,057 Kbp, which is roughly five times higher than the N50 values of the *C. burkhardae* assemblies (148.31 to 460.47 Kbp, Table 1). Additionally, the *H. seosinensis* genome exhibits a notably lower GC content of 46.8% compared to ∼70% in *C. burkhardae* (Table 1), a substantial difference between these related but distinct bicosoecid genera. Like *C. burkhardae* (Hackl et al., 2020), the *H. seosinensis* nuclear genome appears to be diploid, as suggested by a unimodal distribution of the two most frequent alleles in a ploidyNGS analysis (Supplementary Fig. S3b). We mapped six RNA-seq data sets (Harding et al., 2016) to the assembly, with 95% of reads aligning, indicating a high degree of completeness. Overall, the assembly metrics are indicative of a high-quality assembly, and the true completeness may be even higher than estimated, as BUSCO-based analyses tend to underrepresent the gene content of protists (Saary et al., 2020).

**Table 1:** Nuclear genome assembly and annotation statistics for *Cafeteria burkhardae* and *Halocafeteria seosinensis*.

| Assembly | #<br>Contigs | Assembly<br>(Mbp) | Contig N50<br>(Kbp) | % GC | #<br>Proteins | Repeats<br>(%) | BUSCO<br>eukaryote (%) | BUSCO<br>stramenopile (%) |
| --- | --- | --- | --- | --- | --- | --- | --- | --- |
| *Cafeteria burkhardae* |  |  |  |  |  |  |  |  |
| CrE4-10P<sup>1</sup> | 218 | 35.34 | 402.89 | 70.5 | 8364 | 5.1 | 65.9 | 87.7 |
| CrBVI<sup>1</sup> | 170 | 36.33 | 460.47 | 70.1 | 8483 | 8.88 | 65.9 | 87.8 |
| CrCflag<sup>1</sup> | 270 | 34.52 | 231.39 | 70.5 | 8018 | 6.05 | 66.7 | 86.1 |
| CrRCC970<sup>1</sup> | 396 | 33.99 | 148.31 | 70.5 | 7857 | 4.02 | 61.2 | 85.1 |
| *Halocafeteria seosinensis* |  |  |  |  |  |  |  |  |
| HsEHF34 | 60+mito | 38.89 | 1057.34 | 46.8 | 11684 | 19.32 | 78.3 | 81.9 |
| HsEHF34<sup>2</sup> | 8058 | 44.38 | 45.1 | 46.87 | — | — | 77.5 | 82.1 |
| <sup>2</sup>Harding et. al., 2016; <sup>1</sup>Hackl et. al., 2020. |  |  |  |  |  |  |  |  |

Repetitive sequences account for approximately 19% of the *H. seosinensis* assembly (Fig. 1a), substantially higher than the 4–9% observed in the *C. burkhardae* assemblies (Table 1, Supplementary Fig. S3c). Among the repeat categories, LINE retrotransposons are the most abundant type of mobile element identified (∼4%), followed by DNA transposons (2.3%) (Supplementary Fig. S3c). A substantial portion of the repeat fraction could not be classified using standard detection software, consistent with the presence of distinct, uncharacterized mobile elements or fragments thereof.

### Gene content

Protein-coding genes predicted using Braker2 were selected over those generated with Funnanotate due to the higher number of BUSCO orthologs identified. A total of 11,684 such genes were predicted for *H. seosinensis*, substantially higher than the average of 8,180 gene models predicted for the four similarly sized assemblies of *C. burkhardae* (Table 1). This finding aligns with the previously reported high proportion of duplicated genes in *H. seosinensis* (Harding et al., 2016). It is possible that observed differences in gene number and redundancy in part reflect technical limitations related to assembly fragmentation and limited RNA-seq data available for *C. burkhardae*. Nevertheless, given their similar assembly sizes and completeness scores, the *H. seosinensis* and *C. burkhardae* genome annotations are broadly comparable, and provide a suitable framework for exploring gene content variation between the two genera.

### Ortholog inference and enrichment analysis

Orthologous gene family analysis was conducted using the predicted proteomes of the four *C. burkhardae* strains (Hackl et al., 2020) and those inferred here for *H. seosinensis.* From a total set of 42,301 protein-coding genes, 8,572 orthogroups were identified using OrthoFinder v.1.14 (Fig. 1c). Among these, 3,731 orthogroups were found to be shared between both species and all strains (Fig. 1c). Approximately 63% of the gene content of *H. seosinensis* is shared with the four *C. burkhardae* strains, supporting their close phylogenetic relationship (Fig. 1b). To gain broader functional insights, we conducted a gene ontology (GO) enrichment analysis on the *H. seosinensis-*specific gene fraction contained in 910 orthogroups versus the 3,731 core orthogroups shared among the two bicosoecid genera. The analysis revealed several enriched GO terms in the *H. seosinensis* unique gene fraction (versus the *C. burkhardae* core). These enriched terms are broadly associated with four main processes: (1) cellular signaling and stress response (e.g., calcium ion binding and response to biotic stimuli), (2) chromatin modification (e.g., ATP-depending chromatin remodeler activity and histone H3 methyltransferase activity), (3) nucleic acid metabolism (e.g., guanyl nucleotide binding), and (4) mobile elements (e.g., DNA integration) (Fig. 1d). We also identified enriched terms related to protein modification processes, such as phosphorylation and ubiquitination, which may be linked to the adaptation of *H. seosinensis* to hypersaline environments, including the increased hydrophilicity of its cytoplasmic proteome (Harding et al., 2016). Additionally, the enrichment of nucleic acid and chromatin modification categories may reflect fine-tuned regulation of gene expression for critical genes involved in managing osmotic stress, such as ectoine hydroxylase (Harding et al., 2016). At the same time, enrichment of terms associated with the mobilization of TEs (transposable elements), such as DNA integration and aspartyl peptidases encoded by retrotransposons, is likely related to their abundance in the *H. seosinensis* genome. This could also include viral elements such as virophages, given that hundreds of endogenous viral copies have been identified in the *C. burkhardae* genome (Hackl et al., 2021). Overall, our enrichment analysis provides a preliminary glimpse at the unique genetic features of *H. seosinensis*, highlighting key processes potentially linked to high-salinity adaptation and the biology of mobile elements and endogenous viruses.

## Materials and Methods

### Cell culture

*H. seosinensis* strain EHF34 was originally collected and isolated by Park et al. (2006) and maintained at low density in seawater-based media with 150 ppt salinity at room temperature. To isolate *H. seosinensis* cells from prokaryotic contaminants, single-cell picking was performed during the mid-exponential growth phase. Enrichment of single-cell derived cultures was achieved by feeding *H. seosinensis* with the euryarchaeon *Haloferax volcanii,* initially identified as a cobiont (Harding et al., 2016). This culturing strategy facilitated the post-sequencing decontamination process by bioinformatically removing sequences matching the publicly available *H. volcanii* genome sequence (Hartman et al., 2010).

### DNA extraction, genome sequencing and decontamination

Cells were pelleted by centrifugation at 4,000 × *g* for 10 min at 4°C. Genomic DNA was extracted using a combination of the CTAB method (Schenk et al., 2023), phenol-chloroform extraction, and the QIAGEN MagAttract HMW DNA kit. HMW DNA was selected by removing small DNA fragments (<10 Kbp) using the PacBio SRE kit. A genomic library was prepared using the Nanopore ligation kit (SQK-LSK109) and sequenced on a MinION flow cell (FLO-MIN106, R9.4.1 chemistry). Reads were base-called using Guppy v. 3.3 with default parameters (Wick et al., 2019).

The *H. seosinensis* EHF34 genome was assembled using Canu v.2.1 (Koren et al., 2017). The assembly was polished using three rounds of Racon v.1.4.3(Vaser et al., 2017), followed by Nanopolish v.0.8 (Loman et al., 2015). Four additional rounds of correction were performed using Illumina sequence data (BioProject: PRJNA301433) and Pilon v1.23 (Walker et al., 2014). For decontamination, contigs were manually binned based on sequence depth, transcriptomic coverage, GC content, and the taxonomic origin of the best BLAST matches for the open reading frames on each contig. To improve contiguity, the longest reads associated with the eukaryotic fraction of the initial assembly were retrieved using Minimap2 v2.2.17 (Li, 2018) and Filtlong v.0.2 (https://github.com/rrwick/Filtlong) and then reassembled using Canu v.2.1, followed by polishing using Racon v.14.3 and Unicycler (Wick et al., 2017). Haplotypes not resolved in the initial assembly were reconciled using Purge Haplotigs v1.1.2 (Roach et al., 2018). Assembly completeness was assessed using Benchmarking Universal Single-Copy Orthologs BUSCO v.5.2.2 (Simão et al., 2015), using the eukaryota_odb12 reference set (Tegenfeldt et al., 2025). The final assembly consisted of 60 contigs plus an additional contig corresponding to the mitochondrial genome (Supplementary Figure 1).

### Gene prediction, repeat content and ploidy

The mitochondrial genome was manually identified and separated from the nuclear genome assembly (Supplementary Figure 1B). Repeat elements were detected and classified using RepeatModeler (Flynn et al., 2020) and subsequently masked from the genome using RepeatMasker prior to protein-coding gene prediction. Further repeat identification and annotations were performed using the Extensive de-novo TE Annotator (EDTA) (Ou et al., 2019; Su et al., 2021). Gene models were inferred using two *ab initio* pipelines: Braker2 v2.1.5 (Brůna et al., 2021) and Funannotate v.1.8.15 (https://github.com/nextgenusfs/funannotate). Both pipelines utilized GeneMark-ET (Ter-Hovhannisyan et al., 2008) and Augustus (Stanke and Morgenstern, 2005) to generate gene models, supported by spliced-aware mapping of RNA-seq reads using Hisat2 v2.2.1 (Kim et al., 2015). The six *H. seosinensis* RNA-seq libraries used in our analysis were obtained from NCBI: SRX1423011, SRX1423005, SRX1423004, SRX1421934, SRX1423015, SRX1423016 (Harding et al., 2016).

Protein annotations were conducted with Funannotate using UniProtDB v.2022_03, EggNog v. 1.0.3 (Huerta-Cepas et al., 2017), MEROPS v.12.0 (Rawlings et al., 2018), CAZYme, and PFAM-A 35.0 (Finn et al., 2014) with an e-value cut-off of 1E-3. Ploidy was inferred using NGSploidy v.3.1.3 (min_cov =10) (Augusto Corrêa dos Santos et al., 2017).

The *H. seosinensis* mitochondrial genome was annotated using the MITOS2 automated annotator (Bernt et al., 2013). Genome visualization was performed using OrganellarGenomeDRAW (OGDRAW) (Lohse et al., 2013).

### Comparative genomics and phylogenetic analysis

A pangenome analysis was performed using *Halocafeteria seosinensis* and four strains of *Cafeteria burkhardae* (assemblies: CrBVI, GCA_008330645.1; CrE410P, GCA_008330665.1; CrCflag, GCA_008330625.1; CrRCC970, GCA_008330635.1) (Hackl et al., 2020). Repeat content for the four *C. burkhardae* strains and *H. seosinensis* was predicted using RepeatModeler2 and EDTA (Flynn et al., 2020; Ou et al., 2019; Su et al., 2021). Gene clustering was conducted using OrthoFinder v.1.14 (Emms and Kelly, 2015). Enrichment analysis was performed using ClusterProfiler (Wu et al., 2021) on genes associated with the 910 unique orthogroups identified in *H. seosinensis* compared to the core orthogroups shared between four *Cafeteria* strains and *Halocafeteria*.

For 18S rRNA gene phylogenetic analysis, genes were identified in the *H. seoensis* genome using Barrnap (https://github.com/tseemann/barrnap). Additional 18S rRNA sequences from closely related strains, species and genera were obtained from NCBI (*C. burkhardae,* AF174354.1; *Cafeteria.* sp., AF174366.1; *C. atacamiensis* ON412942.1; *C. maldiviensis,* MN315609.1; *H. seosinensis* clone 200, KT210044.1; *H. seosinensis,* DQ269470.1; *H. seosinensis* clone 3, KT210044.3; *Caecitellus paraparvulus*; DQ220713.1; *C. parvulus,* AF174368.2.) (Jeong et al., 2023; Park et al., 2006). The 18S rRNA sequences (∼1,600 sites) were aligned using MAFFT v7.525 (Katoh and Standley, 2013) with subsequent manual curation. Phylogenetic inference was performed using IQ-TREE v. 2.3.6 (Nguyen et al., 2015) with GTR+F+G using 1,000 ultrafast bootstrapping replicates. The tree was rooted using *Caecitellus paraparvulus* and *C. parvulus*.

## Supplementary Material

Supplementary material is available at *Genome Biology and Evolution* online.

## Acknowledgments

We thank Tommy Harding and Alastair Simpson for advice on the culturing of *Halocafeteria seosinensis*.

## Data Availability

The nanopore-based assembly, annotation and Sequence Read Archive (SRA) data for *H. seosinensis* are available on NCBI under PRJNA1135893. SRA: SRR29828113; SRR29828114.

## SUPPLEMENTARY MATERIAL

**Fig. S1.**
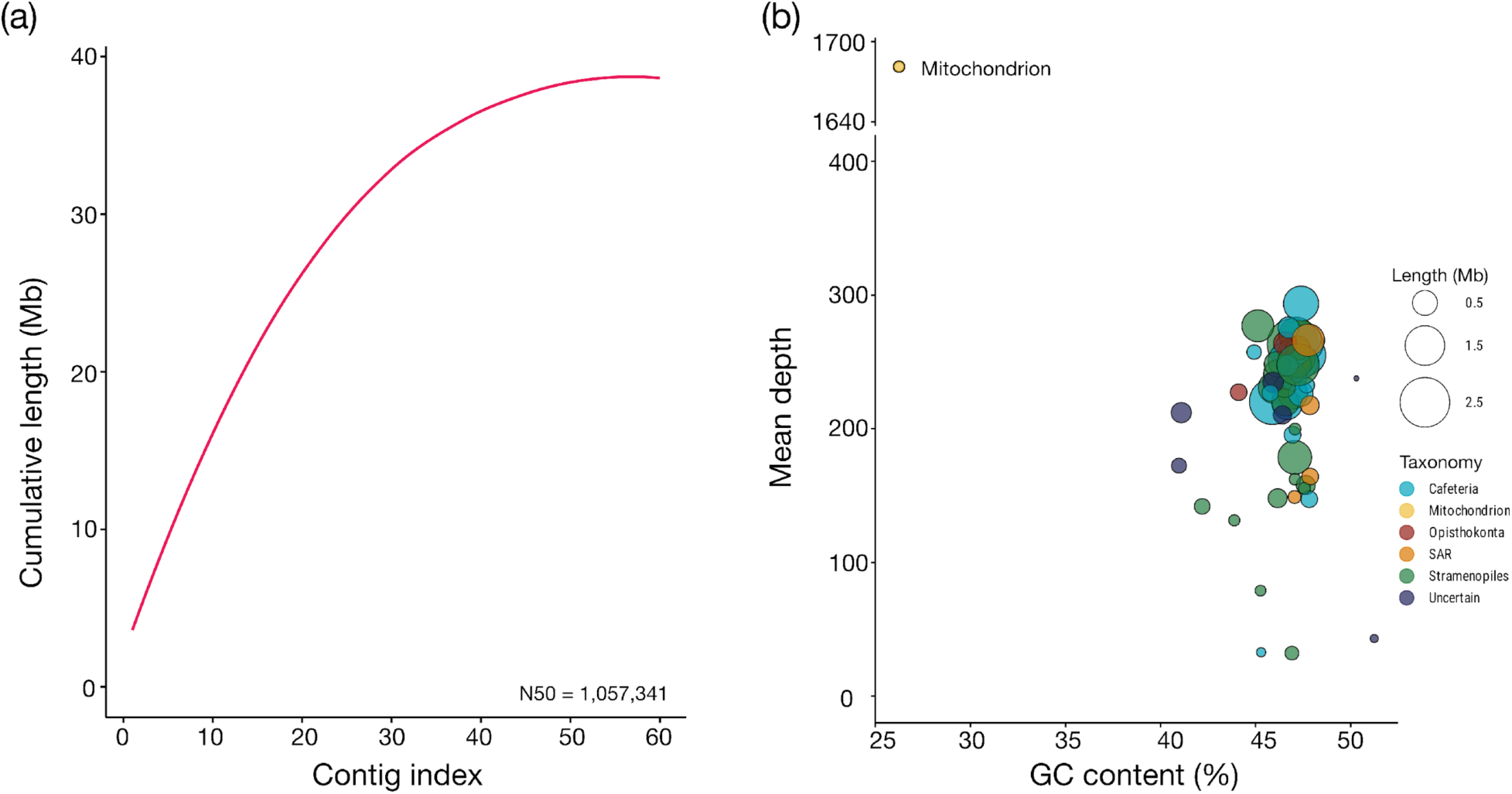
Features of the *Halocafeteria seosinensis* genome assembly. a) Cumulative sequence length plot. b) Blob plot of *H. seosinensis* contigs filtered based on taxonomic annotation, GC content, and sequencing depth. Each contig sequence is represented as a circle sized according to its length in megabases (Mb) and mapped based on GC content (x-axis) and sequencing depth (y-axis).

**Fig. S2.**
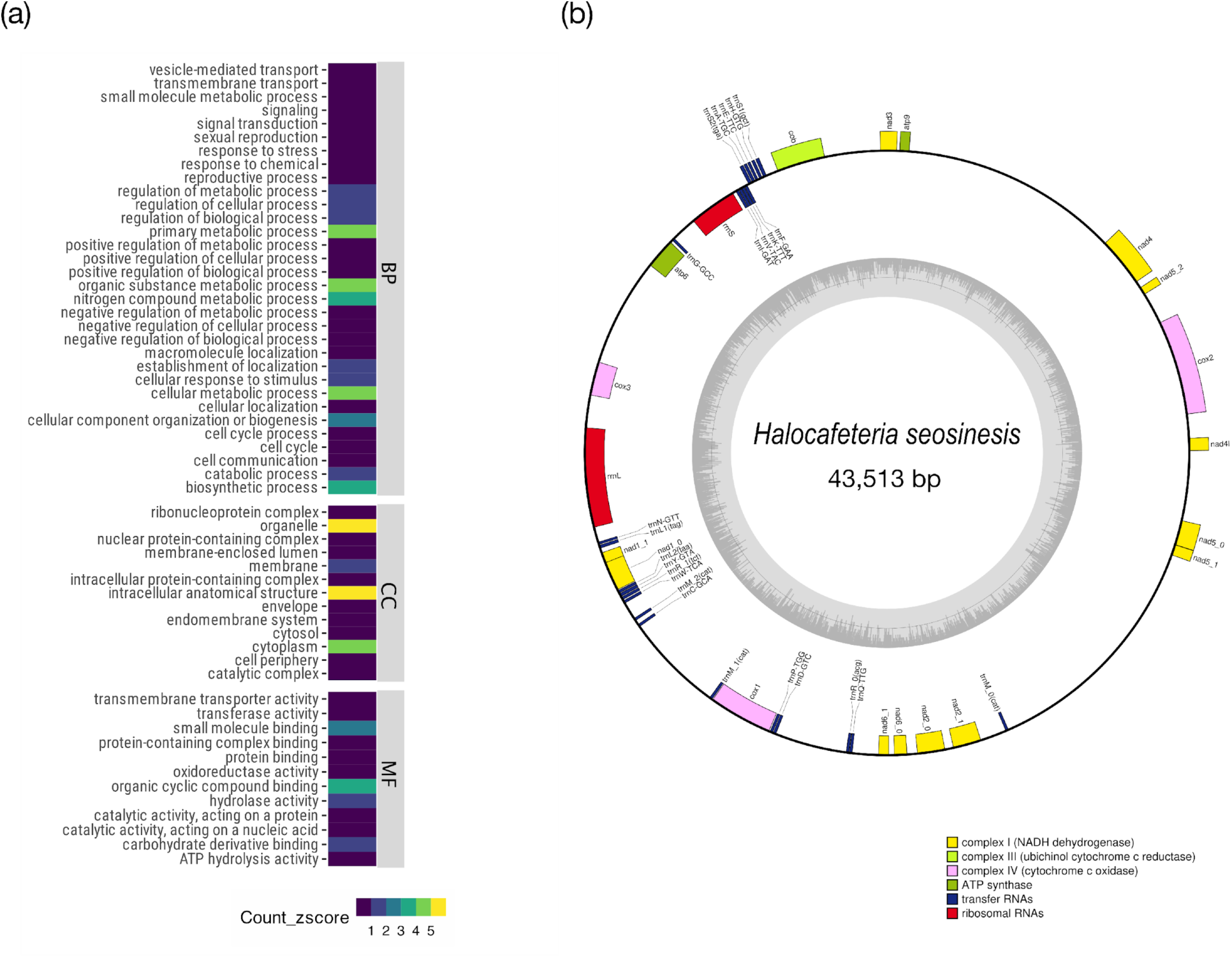
Gene ontology annotations of the *Halocafeteria seosinensis* nuclear and mitochondrial genomes. a) Nuclear gene ontology annotation of the core protein-coding genes associated with the 3,606 orthogroups shared between *H. seosinensis* and four strains of *Cafeteria burkhardae*. BP, CC and MF correspond to biological processes, cellular components, and molecular functions, respectively. The most prevalent categories (z-score>0) are shown. b) Physical map of the ∼43 Kbp mitochondrial genome of *H. seosinensis*; functions of the predicted genes are shown.

**Fig. S3.**
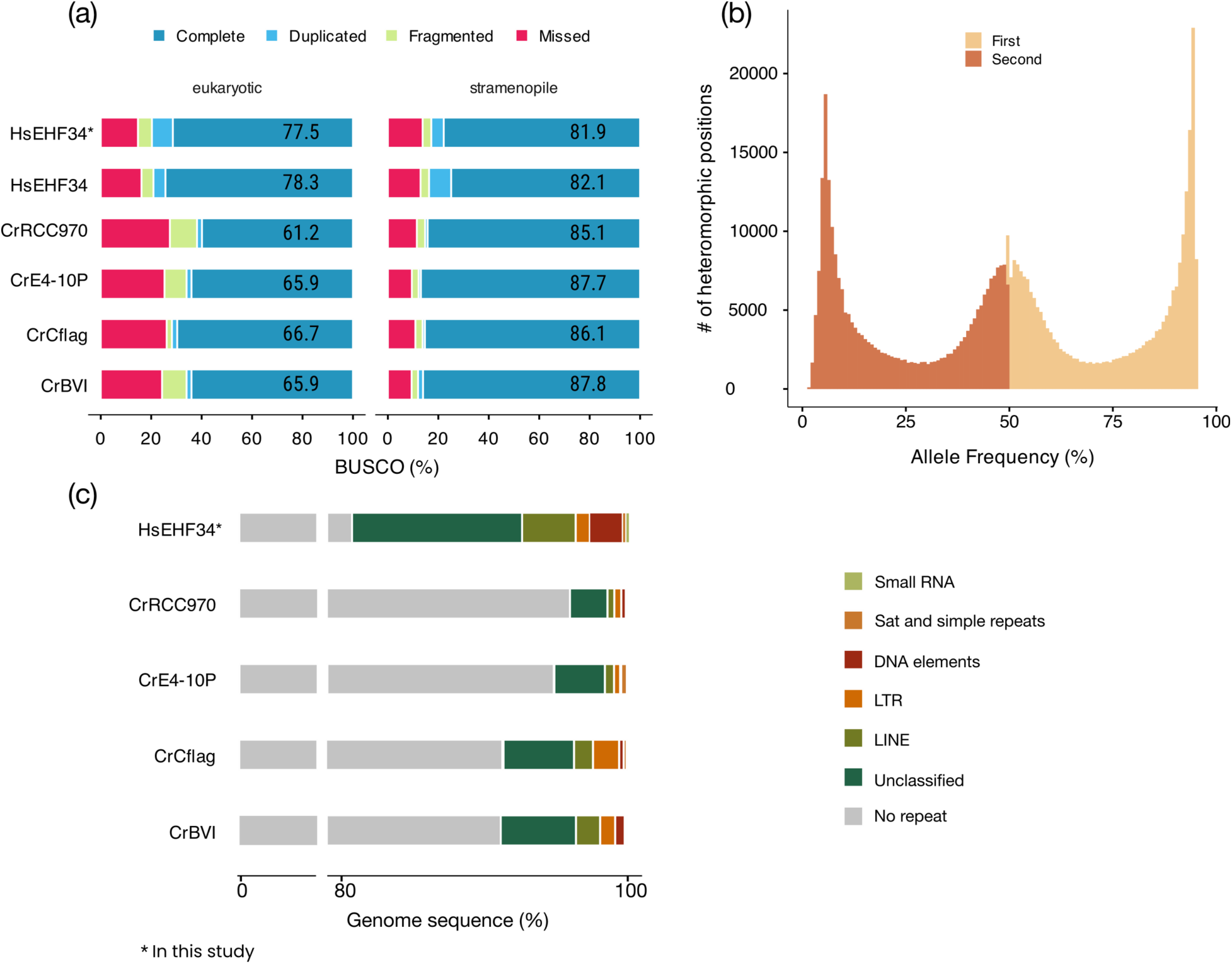
Nuclear genomic features of *Halocafeteria seosinensis* strain EHF34. a) Comparative assessment of genome completeness based on BUSCO orthologs (BUSCO v.5.2.2, odb12) and predicted proteomes of the short read (HsEHF34) and long read (HsEHF34*) assemblies of *H. seosinensis*, as well as four *C. burkhardae* (Cr) strains (Hackl et al., 2020). BUSCO ortholog categories are color-coded. b) Histogram showing unimodal distribution of the two most common alleles (i.e., first and second), suggesting a diploid genome for *H. seosinensis*. Ploidy levels were determined using ploidyNGS (v.3.1.3). c) Comparison of repeat content among four strains of *C. burkhardae* and *H. seosinensis,* highlighting the most prominent groups of repetitive elements. Genomic regions without obvious repeats were classified as “No repeat”, while “Unclassified” refers to repeats with no detectable similarity to repeat sequences in the RepeatMasker library (RepeatMasker v.4.1.1).

